# Coarse-grained simulations of long intrinsically disordered proteins: a benchmark of Martini 3 force-fields

**DOI:** 10.64898/2026.07.17.739185

**Authors:** Christina Goss, Camilo Aponte-Santamaría, Frauke Gräter

## Abstract

Martini 3 is a force field ideally suited to simulating long intrinsically disordered proteins (IDPs) in cell-like surroundings. So far, most Martini 3 variations intended for IDPs have only been benchmarked on shorter IDPs of up to 140 amino acids. In this paper, we present a comprehensive benchmark including IDPs up to 809 amino acids in length and compare the behavior of four well-known Martini 3 variations for IDPs. Modifications to only the bonded parameters result in excessively compact conformations, thereby failing to reproduce the experimental radius of gyration observed for large IDPs. In contrast, general rescaling of interaction parameters, including tuning electrostatic interactions in the case of highly-charged long IDPs, yields acceptable levels of compaction at all tested length scales.

## Introduction

Intrinsically disorder proteins (IDPs) are proteins that do not fold to a single well-defined structure but flexibly shift through a wide range of conformations. Because typical structure-based methods do not work straightforwardly on them, it is particularly difficult to establish a relation between their structure and function. One promising computational method is to study their dynamics and conformational ensembles through molecular dynamics (MD) simulations [1].

Creating force fields that accurately capture the dynamics of IDPs has historically been fraught with difficulties. Force fields are often optimized for folded proteins, and this can lead to overly compact conformations when used for intrinsically disordered proteins [1]. Slight corrections to protein or water properties have resulted in well-established all-atom force fields, such as Charmm36m [2] or the TIP4PD water model [3], that can successfully simulate both folded and disordered proteins. Nevertheless, due to the computational cost involved in atomistic MD simulations, these models have been mainly restricted to IDPs of moderate size, benchmarks typically simulating nothing larger than *α*-synuclein (140 amino acids) [3, 4].

Alternatively, coarse-grained MD simulations, where groups of atoms are represented by single beads, allow for more efficient sampling of the conformational dynamics of even larger IDPs. Forcefields such as HPS [5, 6], CALVADOS [7, 8] or MPiPi [9] have been tuned specifically to replicate the properties of single-chain and condensates of IDPs. These models are very coarse-grained, using one bead per amino acid and implicit water, allowing for efficient MD simulations. In fact, such models have recently enabled high-throughput MD simulations of the entire intrinsically disordered proteome, including both short and long disordered fragments [10]. However, this improvement in performance comes at the expense of a loss in chemical detail.

One of the most versatile coarse-grained force fields for biomolecular systems is Martini 3 [11]. It spans lipid membranes [12], proteins, explicit water, a wide variety of ligands and is ever expanding [13]. Martini 3 maps two - four non-hydrogen atoms onto one coarse-grained bead, preserving to a good extent chemical detail while speeding up simulations considerably. While Martini 3 has been used to successfully study a wide range of systems, from complex lipid bilayers to synthetic polymer systems [14], it is however known to underestimate the global dimensions of IDPs [15].

A variety of solutions have been proposed to successfully model IDPs with Martini. First, Thomasen et al. demonstrated that increasing the protein-water interaction (later referred to as increase P-W), by rescaling the *ϵ* parameter of each Lennard-Jones potential by 1.10, led to more accurate predictions of the global dimensions of IDPs [16]. However, this increase changes the balance of interactions in more complex systems that feature more than IDP-water interactions, such as protein-membrane systems. Thomasen et al. proposed that the same improvement in predicting the size of IDPs, was also possible by decreasing the protein-protein interaction by 0.88 [17] (later referred to as decrease P-P), without disturbing the balance of protein-membrane interactions.

Two other methods have addressed the problem of overly compact IDPs without rescaling the non-bonded interactions. Firstly, in the GōMartini 3 forcefield a Gō-type virtual bead that interacts only with water was added to the backbone beads of IDPs. This achieved the same effect of increasing protein-water interactions, as the first method (increase P-W), but without having to rescale any of the interactions between Martini beads [18]. Secondly, instead of tuning the non-bonded interactions, the bond distances, angles and dihedrals were re-parameterized to better match all-atom IDP simulations, resulting in the promising Martini3-IDP forcefield [15] which has been conveniently integrated into the Martini ecosystem.

All four of these methods greatly improved the agreement between the radius of gyration (*R*_*g*_) obtained by simulations and that measured by small-angle X-ray scattering (SAXS). However, the test sets for each method consisted of small to medium sized IDPs. The Martini3-IDP forcefield, in particular, only considered IDPs up to 140 amino acids in length. IDPs of several hundreds of amino acids are also found in proteomes and mediate key biological processes. For instance, the inner centriole protein (INCENP) contains a ∼440 AA disordered region which plays a key role during cell division [19], CALM and its neuronal specific homolog AP180 contain disordered regions ∼ 600 amino acids in length necessary for clathrin coated vesicle formation [20] and the 800 amino long disordered region of Lubricin lubricates synovial joits [21]. The great promise of the Martini forcefield is that with the speed gained by coarse-graining, the larger time- and length-scales inherently required for ultra-long IDPs become accessible.

Here, we have tested each proposed method of simulating IDPs using Martini 3 on a test set including IDPs up to 800 amino acids long. All methods predict the compactness of IDPs of lengths below 140 amino acids reasonably well. Nevertheless, we find that Martini 3 IDP and the GōMartini method fail to correctly estimate the radius of gyration of large IDPs, while the two scaling approaches prevail. To further explore what is necessary to simulate large IDPs, we also considered the recent Martini3-NMR method [22, 23], which, though not explicitly developed for IDPs, did well when applied to IDPs in the coarse-grained model CALVADOS [23]. Finally we assessed the effect of tuning the electrostatic interactions, as we have recently shown this to improve the conformation and aggregation of another long disordered biopolymer, namely sugar chains [24]. Based on an analysis of the self-interactions of the IDPs we attribute the overly compact conformations to overall slightly elevated interactions and strong charge-interactions, to which rescaling of interactions is the current best solution.

## Results and discussion

### Several Martini 3 IDP methods predict the size (in *R*_*g*_) of large IDPs poorly

To create our test set, we considered IDPs whose *R*_*g*_ had been experimentally determined by SAXS, as has been done previously for each proposed method. In addition, we considered large IDPs for which the *R*_*g*_ value has been determined either by small angle neutron scattering (SANS) [25], or computed from hydrodynamic radii determined by fluorescence correlation spectroscopy (FCS) [26, 27]. In the latter case, for ideal Gaussian chains, the hydrodynamic radius is related to the radius of gyration by the ratio *R*_*g*_*/R*_*h*_ = 1.24 [28], which Pesce et al. confirmed was approximately 1.2 for the larger IDPs in their experimental data set [29]. For simplicity, we have used the factor of 1.2 to convert the *R*_*h*_ values into *R*_*g*_ values. In total, our test set consists of 11 IDPs, ranging from 24 to 809 amino acids in length and having a *R*_*g*_ between 1.4 and 9nm (see Table 1).

**Table 1:** Test set of IDPs considered in this study.

| Protein | Number of amino acids | $R_g$ [nm] | T [K] | $c_s$ [M] | pH | Source |
| --- | --- | --- | --- | --- | --- | --- |
| Hst5 | 24 | $1.38 \pm 0.14$ | 293 | 0.15 | 7.5 | Jephthah et al. [30] |
| ACTR | 71 | $2.65 \pm 0.3$ | 278 | 0.2 | 7.4 | Kjaergaard et al. [31] |
| A1 | 135 | $2.61 \pm 0.11$ | 296 | 0.05 | 7.0 | Bremer et al.[32] |
| α-synuclein | 140 | $3.55 \pm 0.04$ | 293 | 0.2 | 7.4 | Ahmed et al.[33] |
| K25 | 185 | $4.1 \pm 0.2$ | 288 | 0.15 | 7.4 | Mylonas et al.[34] |
| PNt | 334 | $5.13 \pm 0.01$ | 298 | 0.15 | 7.5 | Riback et al.[35] |
| GHR-ICD | 351 | $6.0 \pm 0.5$ | 298 | 0.35 | 7.3 | Pesce et al.[29] |
| ht40 | 441 | $6.5 \pm 0.3$ | 288 | 0.15 | 7.4 | Mylonas et al.[34] |
| AP180 | 593 | $7.38 \pm 0.55$ <sup>a</sup> | 298 | 0.15 | 7.4 | Zeno et al.[26] |
| OMM64 | 628 | $9 \pm 1$ <sup>b</sup> | 298 | 0.1 | 7.5 | Poznar et al.[25] |
| GW128SD | 809 | $7.92 \pm 0.72$ <sup>a</sup> | 298 | 0.15 | 8.0 | Waszkiewicz et al.[27] |

<sup>a</sup>Radius of gyration calculated from hydrodynamic radius <sup>b</sup>Radius of gyration measured by SANS

We simulated each IDP from our dataset using either plain Martini 3, one of the four proposed solutions for IDPs: Martini3-IDP, which improved bonded parameters [15]; GōMartini 3, which added a virtual bead to each protein backbone bead that only interacts with water [18]; decreasing protein-protein interactions by 0.88 [17]; or increasing protein-water interactions by 1.10 [16]), or a further method chosen for comparison (Figure 1A). The radius of gyration was determined over a cumulative simulation time of 8*µs*. It is clear that many Martini 3 variations predict the radius of gyration of large IDPs (> 140 amino acids) poorly (Figure 1B). In the following, we compared and analyzed the different test systems in detail.

**Figure 1:**
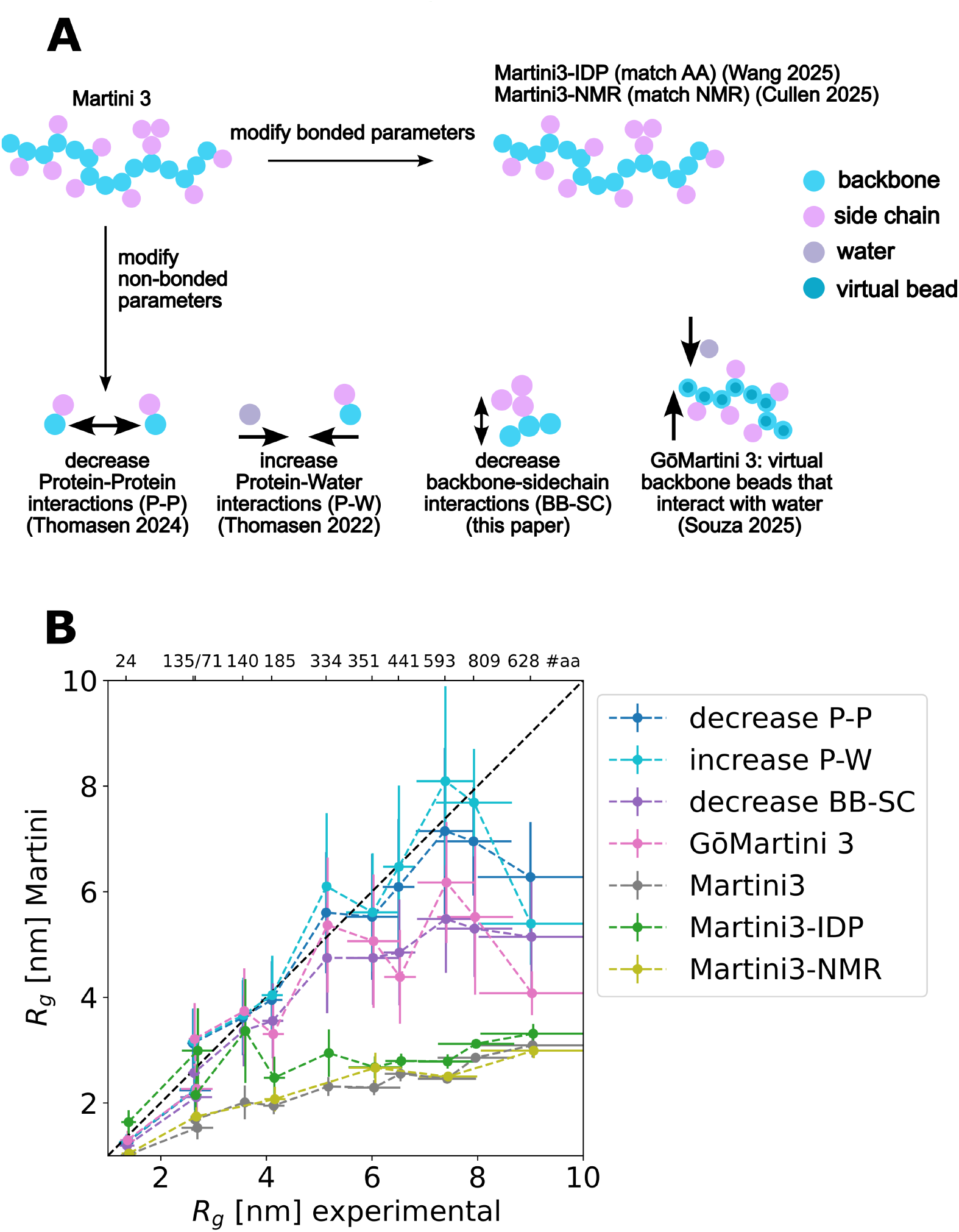
Not all methods for simulating IDPs with Martini 3 describe the size of large IDPs of more than 140 aa. A) The scheme depicts the different Martini 3 variations used to simulate IDPs in this paper. They can broadly be classified to either modify the bonded parameters (top row) or the non-bonded parameters (bottom row). B) The Experimental *R*_*g*_ is plotted against the *R*_*g*_ calculated from simulations with variously modified Martini 3 versions. The length in number of amino acids is indicated at the top. The error bars for the simulations are the standard deviations calculated over 8*µs* of cumulative simulation time.

The different variations of Martini 3 used to simulate IDPs in this paper can be broadly classified to modify the bonded or non-bonded parameters of IDPs (Figure 1 A).

Modifying the bonded parameters of the Martini 3 IDP model is the solution most compatible with the rest of the Martini 3 ecosystem [15]. Martini3-IDPs redefine the bonded terms of the IDP model to better match all-atom simulations and thus improve the prediction of the *R*_*g*_ along with other properties for IDPs up to 140 amino acids in length. An alternative method which can also be classified into this category is Martini3-NMR. This method was recently developed to improve the alignment of Martini 3 simulations with NMR observables [22]. Martini3-NMR helps to preserve the secondary structure of folded Proteins in Martini 3, but it has also showed promise at reproducing the dynamics for the disordered parts of these proteins. In fact, an extension of this model to the coarse-grain forcefield CALVADOS predicted features that were in excellent agreement with experimental measures of IDPs [23]. When comparing the ability of these models to capture the conformational dynamics of IDPs (Figure 1)B), Martini3-NMR offered no improvement on plain Martini 3 when it comes to the tendency for IDPs to take on overly compact conformations. In contrast, Martini3-IDP showed a big improvement for IDPs up to 140 amino acids in length (such as *α*-synuclein, *R*_*g*_ = 3.55 ± 0.04). Nevertheless, the larger the IDP, the more similar the predicted *R*_*g*_ of Martini3-IDP becomes to that of Martini 3 and Martini3-NMR (Figures 1)B and 2). Both methods have a mean absolute error comparable to that of plain Martini 3 (see Figure S1), although Martini3-IDP has slightly better accuracy due to the excellent simulation of shorter (< 140 amino acid) IDPs.

The other variations of Martini 3 modify the non-bonded parameters to better describe IDPs. We now focused on the models that only modified interactions with the protein backbone. GōMartini 3 attempts to disturb the Martini 3 ecosystem minimally by adding a Gō-type virtual bead that interacts only with water to the backbone beads of IDPs, effectively increasing protein-water interactions, and thereby the solubility, without rescaling any other interaction between Martini beads [18]. As an additional possibility, we also applied the decreased Protein-Protein interaction scheme from Thomasen et al. [17], but in this case only to the interactions of backbones with sidechains (labeled as decrease BB-SC). These two models behaved very similarly: IDPs up to 334 amino acids in length took on more extended conformations and thus matched the experimental *R*_*g*_ values (Figure 1B). Above this length, IDPs adopted more compact conformations than expected according to experimental data, but the structures continued to be much more extended than those generated with Martini 3 or variations which modified bonded parameters (compare *R*_*r*_ estimates for “decrease BB-SC” and “GōMartini 3” with estimates for the bond-modifying schemes “Martini3-IDP” and “Martini3-NMR”, in Figures 1B and 2). This is also reflected in the drastic decrease in mean absolute error these methods show in comparison to Martini 3 (see Figure S1).

Finally, the Lindorff-Larsen and Vanni groups have produced two variations of Martini 3 to accurately simulate IDPs: either enhance the solubility by increasing all Protein-Water interactions [16] or make the proteins less “sticky” by decreasing all Protein-Protein interactions [17]. Both of these methods performed well on almost all IDPs of our test set, only underestimating the size of the largest IDP, OMM64 (Figures 1, 2). These two scaling schemes also showed the best mean absolute error of any method in the test set, with decreasing protein-protein interactions displaying slightly better estimates than increasing protein-water interactions (Figure S1).

**Figure 2:**
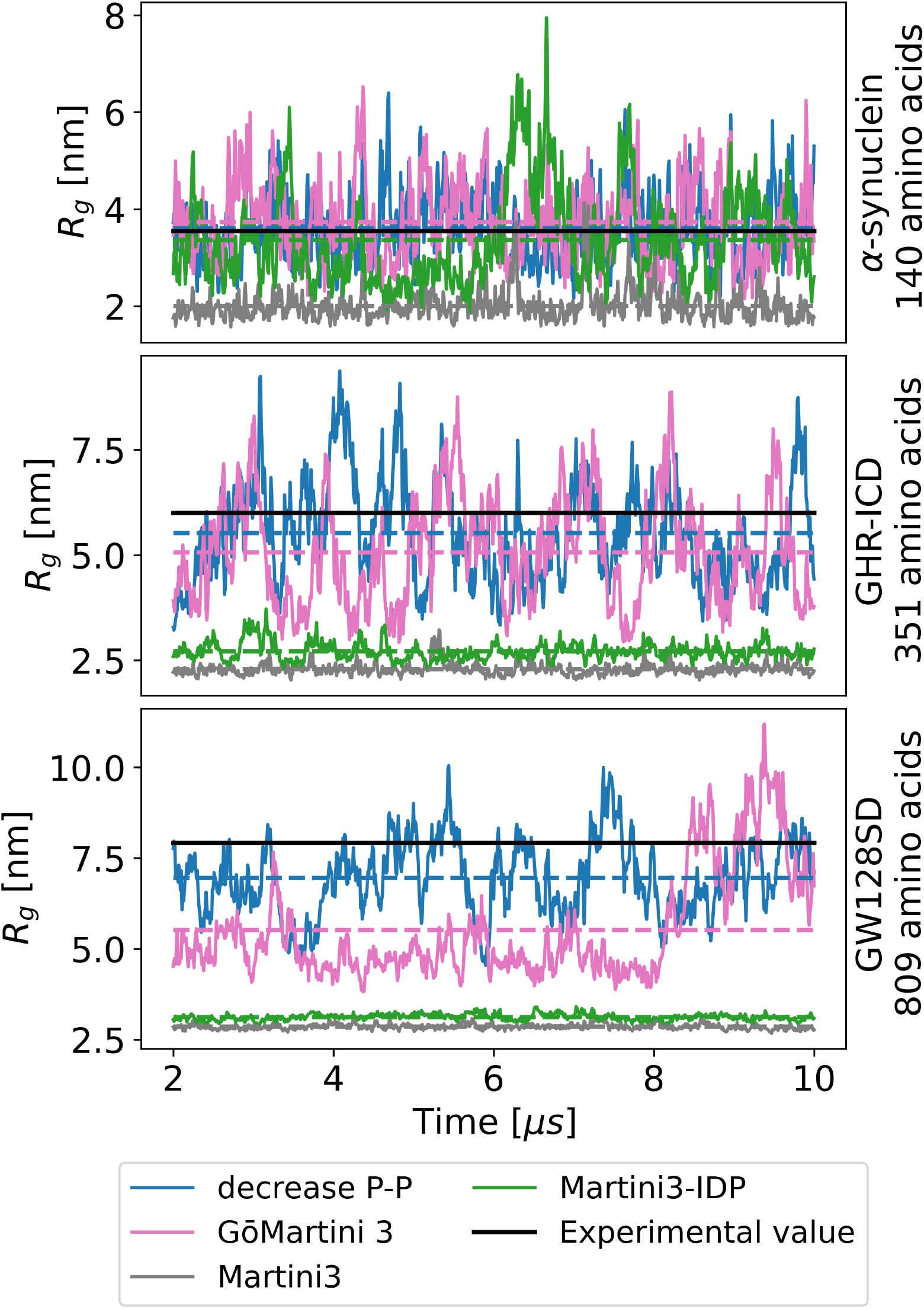
As IDPs get longer than 140 aa, more methods predict the *R*_*g*_ poorly. For smaller IDPs, e.g. *α*-synuclein, all methods inteded to simulate IDPs approach the experimental value. For slightly larger IDPs, such as GHR-ICD, Martini3-IDP shows a more condensed state, while GōMartini 3 and a rescaling Method still approach the experimental value. For the largest of IDPs, here demonstrated on the longest IDP in the test set, GW128SD, only the rescaling Methods approach the experimental values. For further comparison see Figures S2-S5)

### There is no inherent difference in the polymeric properties between small and large IDPs in the studied test set

To understand if a difference in the intrinsic sequence properties of the IDPs themselves causes some simulation methods to produce overly-compact conformations, we calculated a variety of measures that characterize the physico-chemical properties of polymers and are thus used to predict if IDPs will take on compact or expanded conformations (Figure 3). We split our test set into two groups, “small” IDPs ≤ 140 amino acids, for which all schemes to simulate IDPs using Martini predict the *R*_*g*_ accurately, and “large” IDPs > 140 amino acids, where there is considerable variation in how well the Martini variations predict experimental *R*_*g*_.

**Figure 3:**
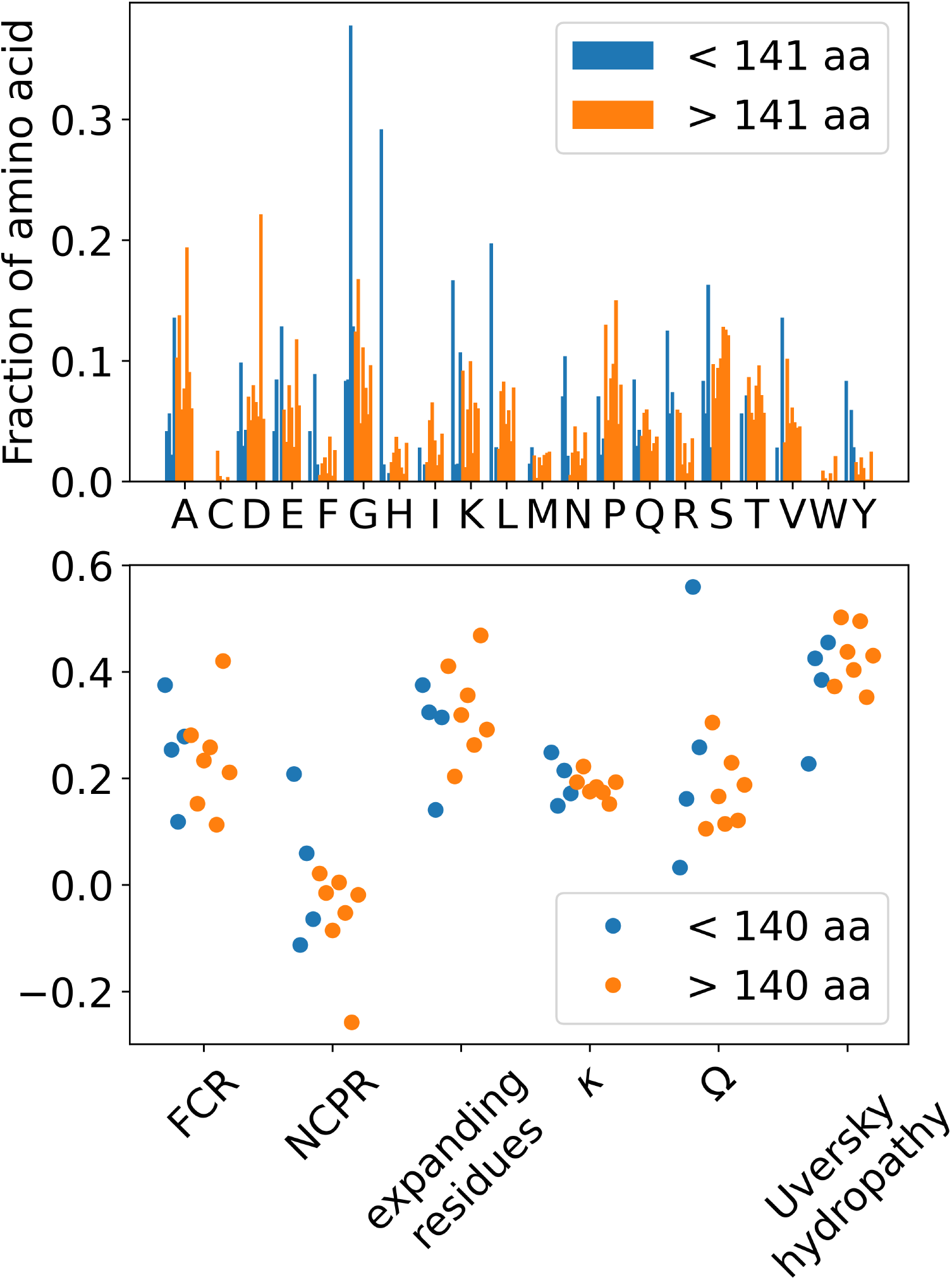
There is no difference in properties known to influence IDP compaction between small IDPs (<141 amino acids, in blue) and large IDPs (>141 amino acids, in orange). Above: The fraction of each IDP that an amino acid is found within. Below: The fraction of charged residues (FCR), Net Charge per Residue (NCPR), fraction of expansion promoting amino acids (R, K, D, E, and P), *κ* parameter, Ω parameter, and Uversky hydropathy is calculated for each IDP in the dataset.

Highly charged IDPs are known to be more extended than those less highly charged [36]. To measure this, we calculated the fraction of charged residues (FCR) and the net charge per residue (NCPR). Neither metric could separate the two size-groups, either visually (Figure 3) or by a Kolmogorov-Smirnov test (Table S1). Beyond the charged amino acids (Arginine (R), Lysine (K), Aspartic Acid (D) and Glutamic Acid (E)), Proline is known to promote more expanded conformations, too [37]. We calculated the fraction of these five expansion promoting amino acids (labeled “expanding residues”) and the fraction each amino acid makes up of the IDP. While there are some differences between groups at the individual amino acid level, the collective proportion of these type of amino acids did not differentiate the two size groups from each other. Beyond the simple composition of the IDP, the arrangement of charged amino acids can also make a difference. The *κ* parameter attempts to measure how clustered the charges are: high values of *κ* (>0.2) indicate likely long-range electrostatic attractions between blocks of oppositely charged amino acids and therefore more compact IDPs [38]. The parameter Ω is an analogous parameter to *κ*, including Proline, too [39]. The two size groups could not be separated from each other by these two parameters either. Finally, the hydrophobicity of a protein can influence its conformation – thus we calculated the Uversky hydrophobicity for all IDPs [40], but were once again unable to use it to seperate the short from the long IDPs in our test set. In summary, the smaller IDPs – for which all proposed methods for simulating IDPs with Martini function well – cover a similar range as the larger IDPs on each metric known to predict the expansion or compaction of IDPs. From this we conclude that, rather than any sequence-specific property, it is the size of the IDPs themselves that cause such a difference in conformations.

### Martini 3 shows strong non-specific interactions, which can be improved by general rescaling

To understand why some corrections result in compact conformations for large IDPs in simulations, and others do not, we inspected the formation of contacts between amino acids of the IDP chains. The average number of non-trivial contacts that each amino acid established, i.e. not with neighboring amino acids, separated the methods that only modified bonded parameters from those that modified non-bonded (Figure 4, above). Martini3-IDP and Martini 3 showed many more distant contacts between residues than the methods that modified non-bonded forces, especially for large IDPs. Furthermore, the methods that modified only the backbone of the protein (GōMartini 3 and decreased BB-SC interactions) displayed a small increase compared to the methods that modified the whole protein (decrease P-P and increase P-W).

**Figure 4:**
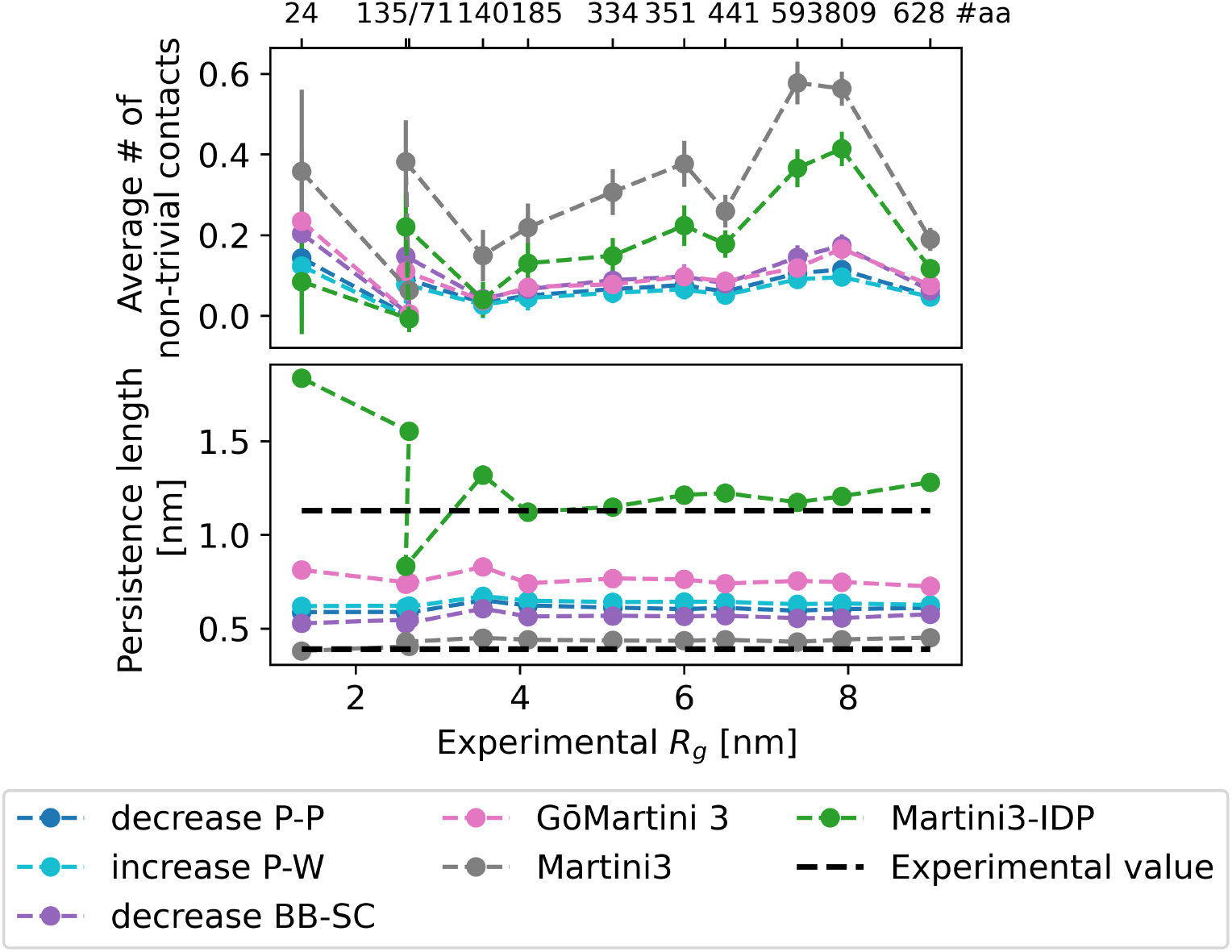
Inter-residue contacts and stiffness recovered for the different Martini schemes proposed for IDPs. Above: The average number of non-neighbor contacts each amino acid has. Martini3-IDP shows pronouncedly more contacts in the dense conformations it takes on for large IDPs. The error bar shows the standard deviation. Below: The persistence length for each simulation method and IDP is plotted against the experimental *R*_*g*_. Martini3-IDP substantially increases the persistence length of the IDP. A range of experimentally measured values is shown in black [41, 42]

A less direct way to measure the interactions between amino acids is to calculate the apparent Flory exponent *ν* [43]. The Flory exponent measures how fast the size of a polymer scales with its chain length. In the limit of polymers in good solvents, *ν* = 0.588 [43], which is close to how the methods that modified the whole protein scale (decrease P-P and increase P-W) (Figure S6). However, when the amino acids interact more strongly with each other, *ν* is lower and the size of IDP chains does not scale as expected with longer and longer IDPs. This is particularly apparent for plain Martini 3 and Martini3-IDP.

We take this as an indication that the Lennard-Jones forces between amino-acids are too large compared to the Lennard Jones forces between amino-acids and water, leading large IDPs to eventually interact too strongly with themselves and form overly compact conformations. While for many IDPs, GōMartini 3 and Martini 3 with decreased backbone-side chain interactions yielded *R*_*g*_ values in good agreement with experiment, for the largest IDP, they sampled overly compact conformations (Figure 1). This indicates that the backbone itself causes much of the compaction, as tuning its interactions is enough to ameliorate the compaction; for very large IDPs, tuning only the backbone is not enough – the side-chain interactions add up to too large a value to successfully match experimental *R*_*g*_ in simulations.

We next asked how the stiffness of the backbone, measured by the effective persistence length, couples to the length scale beyond which we see the IDPs collapse (Figure 4, below). Here, we measured persistence length as the distance at which the local correlation between backbone-bond vectors decays. Comparing the persistence length to the length of the IDP can lead to two distinct regimes: When proteins are much shorter than their persistence length, they behave like rods, but when they are are much longer than their persistence length, they can be modeled as ideal chains. The estimated persistence lengths were found to be in the range of values reported experimentally for IDPs [41, 42]. All corrections made to the Martini forcefield predicted persistence lengths above the values obtained from Martini3. However, changes made to the bonded parameters to create Martini3-IDP effectively increased the persistence length and produced much stiffer proteins. This allowed Martini3-IDP to simulate small IDPs well – the stiffer protein counteracts the stronger non-bonded attraction between beads. However, when the proteins became much larger than the persistence length, this local stiffening could not prevent the self-interaction between more distant parts of the protein, leading again to overly globular states, seen in the original Martini 3. Thus, we believe that deliberately further stiffening the protein is a flawed approach, as there will always be a length scale at which an IDP is dominated by ideal chain behavior, and we risk creating unnaturally stiff short proteins.

To diagnose which type of interactions may be too attractive in the Martini force field, we compared the contact maps between the Martini force fields variants studied here and an all-atom method developed specifically for IDPs. The pattern of which amino-acids preferentially interact with which other amino-acids is very consistent across all Martini-methods and the all-atom force field, the major difference being how strong these interactions are (Figure S7, S8). A closer look at the strength of these interactions revealed an interesting picture: the more compacting forcefields, i.e. Martini3 and Martini3-IDP, showed a much higher proportion of amino-acid pairs that are in contact for short time scales (between 1 and 10% of the simulation), while for GōMartini 3 and the rescaling approaches contacts were much longer lived (i.e. more than 10% of the simulation) (Figure 5). This points to the compaction being a more general “stickiness” problem, with slightly over-estimated non-bonded forces causing attraction among almost any pair of amino-acids, even if only briefly. Slightly tuning down this non-bonded interaction, be it directly (by decreasing the protein-protein or backbone-sidechain bead interaction) or indirectly (by increasing the interactions with water instead), seems to tune down these less-frequent contacts and reproduce the experimental data better.

**Figure 5:**
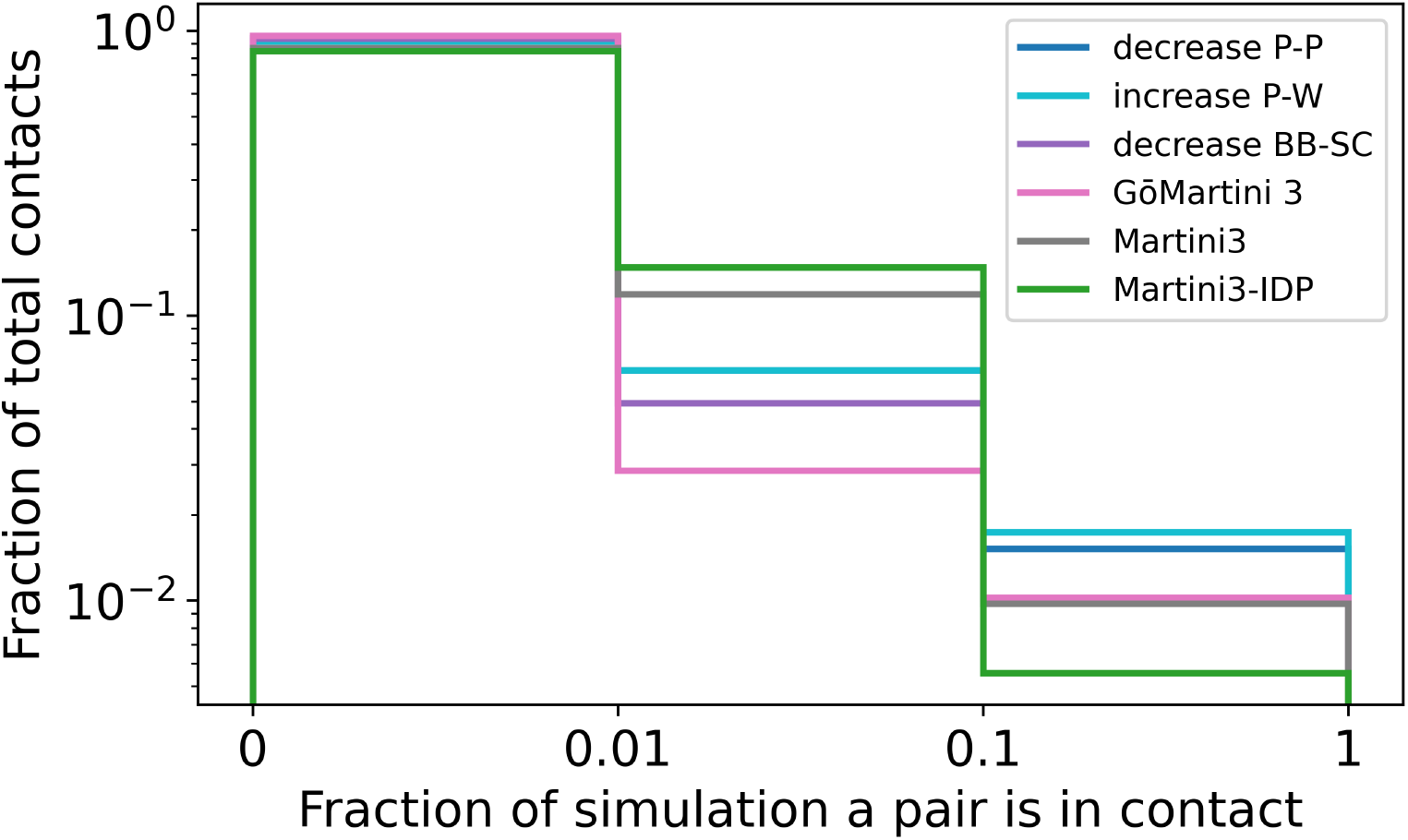
Martini 3 and Martini3-IDP show a high proportion of short contacts (between 1 and 10% of the simulation), while GōMartini 3 and the rescaling methods show a higher proportion of longer, more sustained contacts (more than 10% of the simulation) Histogram of how much of a simulation amino-acids are in contact with each other. The briefest of contacts (less than 1% of the simulation) were very frequent for all Martini variations

One difference between the contact maps of the all-atom force field and the Martini force fields is an increase in interactions for the charged amino-acids (consider for instance Aspartic acid in ACTR or Lysine in *α*-synuclein in Figure S8). This may be caused by ions building salt-bridges where they do not naturally occur, as it has previously been seen for other highly-charged chains parametrized with Martini [24]. Following Greicius et al., we performed some brief experiments modifying the ions (increasing the size and down-scaling the charge to emulate a hydration shell) or modifying the ions and downscaling the charge of the amino-acids to match. When applied to the method which rescales protein-protein interactions [17], modifying the ions resulted in slightly more extended conformations, particularly for OMM64, the largest and most negatively charged IDP in the test set. Less overall rescaling was needed to reproduce the experimental radius of gyration when using modified ions (Figure S10, Figure S11). Surprisingly, using modified ions in combination with GōMartini 3 provided reasonable results on even the largest IDPs, which GōMartini 3 could not successfully simulate on its own (Figure S9). Thus, adjusting electrostatic interactions – in conjunction with the above mentioned corrections – may also help reproducing the compactness of highly-charged large IDP systems.

## Conclusions

Our results show that some of the well known methods for simulating IDPs using Martini 3, although accurate at predicting the conformation of small to medium-sized IDPs, fail at predicting the radius of gyration of large IDPs.

Understanding how and why these methods fail is instructive to create a method that properly samples the conformation of both short and long IDPs. Concerning long IDPs, by improving the bonded parameters, Martini3-IDP effectively increased the persistence length of the IDPs, leading to collapse once the IDPs became about 20 times as long as the persistence length and the attractive forces between beads could not be compensated for any longer. As implemented in GōMartini 3 or the two rescaling approaches, changing the non-bonded forces prevented collapse. GōMartini 3 adds virtual beads only to the peptide backbone, thereby increasing its solvation. For large IDPs, this was not enough to solve the problem of overly compact conformations, but at least the predicted conformations were much more extended than those produced by unmodified Martini 3 or Martini3-IDP. This method could not account for the fact that some of the side-chain beads still interacted too strongly, which is in line with errors in the ranking of dimerization affinity of side chain analogues [16]. When scaling up to IDPs several hundred amino acids long, any slight error can end up having much larger effects. However, in comparing the contact maps between small IDPs simulated with Martini and all-atom force fields, a general “stickiness” was observed, for which a general down-scaling of the non-bonded interaction forces seems to be–at the moment–the best solution. Charged amino-acids show slightly too-strong interactions, which may be mediated by unnatural ion-bridges. A simple solution to this could be to use larger and less charged ions, as suggested in Greicius et al. [24].

In summary, for long IDPs of more than 200 amino acids, our current recommendation is to use either of the rescaling approaches [16, 17]. Depending on the system of interest, decreasing the protein-protein interactions [17] will likely be the best choice, as it does not disturb protein-membrane interactions. A more thorough discussion of the advantages of rescaling protein-protein over protein-water interactions can be found in Thomas et al. [17]. In highly charged systems, we recommend using larger and less charged ions, as suggested in Greicius et al. [24], with a slightly lower rescaling factor of *λ* = 0.92, which gave the best results on our test set.

## Methods

### IDP Simulations

We performed all MD simulations, excepting Martini3-NMR, using GROMACS 2025.3 [44] and the Martini 3 [11], the Martini3-IDP [15], the GōMartini 3 [18] or a rescaled version of the Martini 3 forcefield [16, 17]. Initial conformations were created using Polyply [45] and solvated using GROMACS. Initial box sizes were 25nm x 25nm x 25nm for IDPs shorter than 442 amino acids, and 35nm x 35nm x 35nm for those larger. NaCl concentration was set to match the experimental systems (see Table 1) and neutralize any charges. Energy minimization was performed using steepest descent for 3000 steps. The general simulation protocol used the leap-frog algorithm [46] to integrate the equations of motion. The Verlet neighbor search algorithm was used to update the neighbor list every 20 steps with a cutoff distance of 1.35 nm. Potential shift was used to calculate van der Waals interactions with a 1.1 nm cutoff and Coulomb interactions were treated by a reaction-field with dielectric constant 15 and 1.1 nm cutoff. In the first equilibration step, temperature was brought to match experimental values (see Table 1) using the velocity rescaling thermostat [47] with a coupling constant of 1 ps over 67000 steps with a step size of 15 fs. Following this, the pressure was equlibrated to 1 bar using the Berendsen barostat[48] with an isotropic coupling constant of 4 ps, and a compressibility of 4.5e-5 bar^−1^ over 134000 steps. Production simulations lasted 10 *µs* with a step size of 20 fs and switched to the Parinello-Rahman barostat[49] with a coupling constant of 12 ps. The IDPs were then simulated for 10 *µs* and the first 2 *µs* discarded as equlibaration.

The radius of gyration (*R*_*g*_) was calculated from the trajectories using GROMACS `polystat`. The persistence length was calculated using the python package MDAnalysis [50]. The average number of intra-chain contacts per amino-acid was calculated by applying GROMACS `pairdist` on each amino acid in an IDP. Distances less than 0.4 nm were considered to be contacts, and averaged over time. Trivial contacts (contacts of an amino acid with itself or its neighboring amino acids) were excluded. The apparent Flory exponent *ν* was obtained as the exponent of the fit 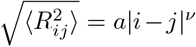, of root-mean-square distance between aminoacids 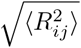 and separation along the linear sequence |*i* − *j*| greater than five residues [10].

### Martini3-NMR

By recommendation of the authors [22, 23], we only simulated IDPs for which s4pred [51] predicted almost exclusively the coil formation (less than 15% were predicted to be non-coil, and no stretches longer than 10). This set consisted of the proteins Hst5 (24 aa), A1 (135 aa), K25 (185 aa), GHR-ICD (351 aa), AP180 (593aa) and OMM64 (628 aa). As there were no complete NMR data sets for these proteins, synthetic chemical shift data were generated using CamCoil [52]. These simulations were performed as Langevin Dynamics simulations in OpenMM 8.2.0 [53] using the Langevin integrator with a time step of 10 fs and friction coefficient of 0.01 ps^−^1, under periodic boundary conditions. The system was set up using GROMACS, as described above. At the beginning of each simulation, the system was energy-minimized by the steepest descent algorithm, followed by 6700 steps of NVT equilibration and 134000 steps of NPT equlibration, with a step size of 15 fs. The step size was increased to 20 fs and the chemical shift (CS) restraints were then gradually imposed by increasing KCS by 0.0005 *kJmol*^−1^ each step until K=25 *kJmol*^−1^ was reached. Finally, 10 *µs* of simulation were performed, and analyzed as above.

### All-atom simulations

The four smallest IDPs of the test set–Hst5, ACTR, A1 and *α*-synuclein– were simulated using all-atom force fields for comparison. The amber99sb-star-ildnp force field was employed for the protein [54, 55] and TIP4P-D for the water molecules [3]. Initial structures were taken from AlphaFold3 predictions [56] were placed within a dodecahedral simulation box with 3 nm between the Protein and the nearest box wall and subsequently they were solvated. Similarly as in the Martini simulations, NaCl concentration was set to match the experimental systems (see Table 1) and neutralize any charges. Energy minimization was performed using steepest descent algorithm until the maximum atomic force was below 1000 kJ mol_1_nm_1_. Temperature was brought to match experimental values (see Table 1) using the velocity rescaling thermostat during 5 ns with a coupling constant of 0.1 ps. Subsequently, the solvent was relaxed in the NPT ensenble at a pressure of 1 bar using the Parinello-Rahman barostat [49] with a coupling constant of 2 ps for 10 ns. Finally, production runs were executed under the NPT ensemble for 500 ns. Analysis was performed as for the Martini simulations.

### Analysis of IDPs

The Analysis of the IDPs was completed using CIDER [57]. A Kolmogorov-Smirnov test was computed between short (< 140 amino acids) and long (> 140 amino acids) for each metric (see Table S1) using SciPy stats [58].

## Acknowledgements

We thank Mina Cullen and Davide Meracadante for their help implementing Martini3-NMR and their fruitful discussions. This research was conducted within the Max Planck School Matter to Life supported by the Dieter Schwarz Foundation and the German Federal Ministry of Education and Research (BMBF) in collaboration with the Max Planck Society. The authors gratefully acknowledge support by the German Research Foundation (DFG) grant GRK 2450. Computations were performed on the HPC system Otter at the Max Planck Computing and Data Facility.

## Supporting information

The following files are available free of charge.

- Suppplement.tex: Further Figures and error analysis consisting of results of Kolmogorov-Smirnov tests (Table S1), error of the test set (Figure S1), time traces of all simulated IDPs (Figure S2-S5), the apparent Flory exponent (Figure S6), contact maps of Martini simulations compared to all-atom simulations (Figure S7-S8), radius of gyration for Martini force-fields when the ions have been rescaled and the corresponding errors (Figure S9-S11).
- Files to generate the simulations as well as analysis Scripts are available on Edmond (https://edmond.mpg.de/)

---

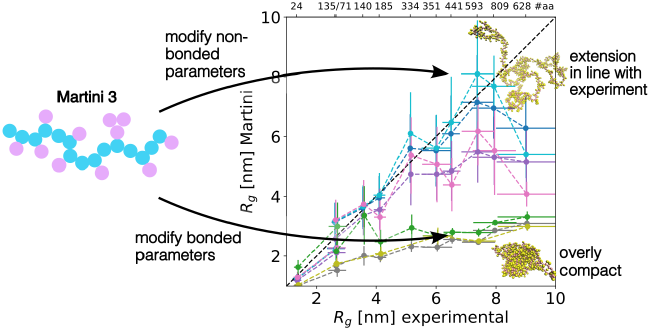

## Notes

### Competing Interest Statement

The authors have declared no competing interest.

