## Supplemental Figures and Table for "Coarse-grained simulations of long intrinsically disordered proteins: a benchmark of Martini 3 force-fields"

July 17, 2026

| Metric | p-value |
|---|---|
| FCR | 0.89 |
| NCPR | 0.42 |
| expanding residues | 0.95 |
| $\kappa$ | 0.78 |
| $\Omega$ | 0.78 |
| Uversky hydropathy | 0.89 |

Table S1: Result of the Kolmogorov-Smirnov test to differentiate between short and long IDPs. No p-value is low enough to reject the null-hypothesis that the listed sequence properties, from both the short and long IDPs, come from the same underlying distribution.

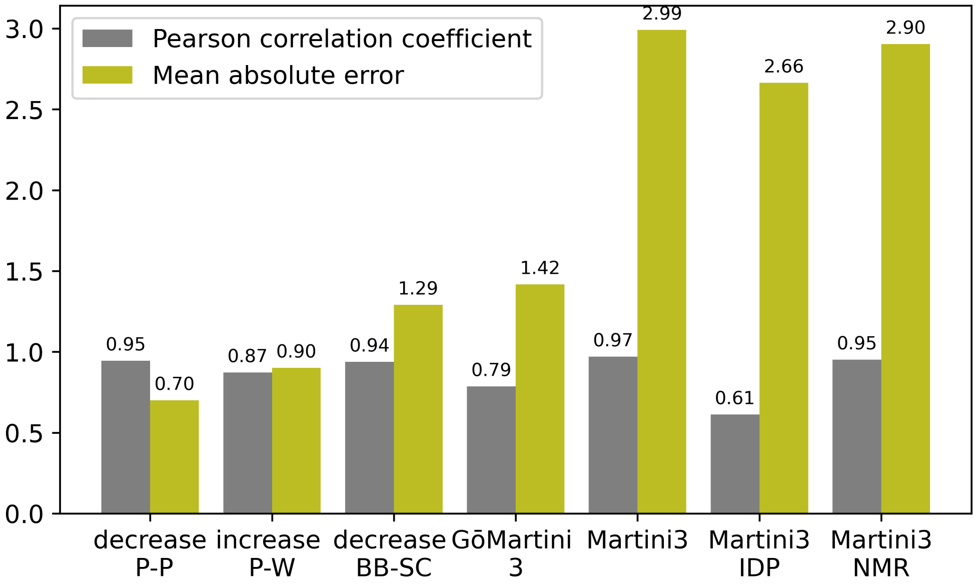


Figure S1: Pearson correlation and mean absolute error on the test set for each Martini 3 variation featured in Figure 1 of main text

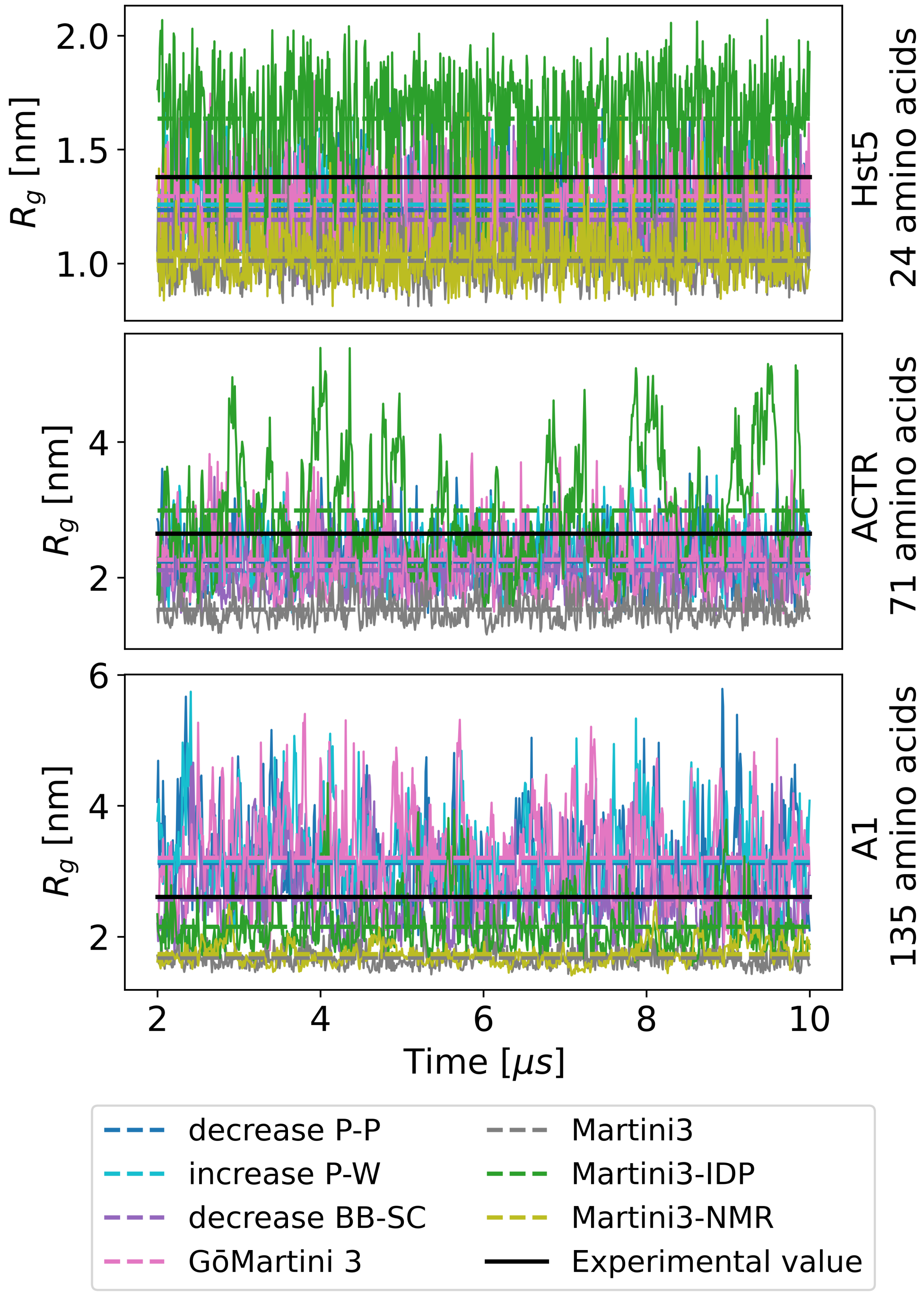


Figure S2: Time traces of the simulation for Hst5, ACTR and A1. Mean values of $R_g$ are shown as dashed lines for each simulation, and the experimental value as a solid black line.

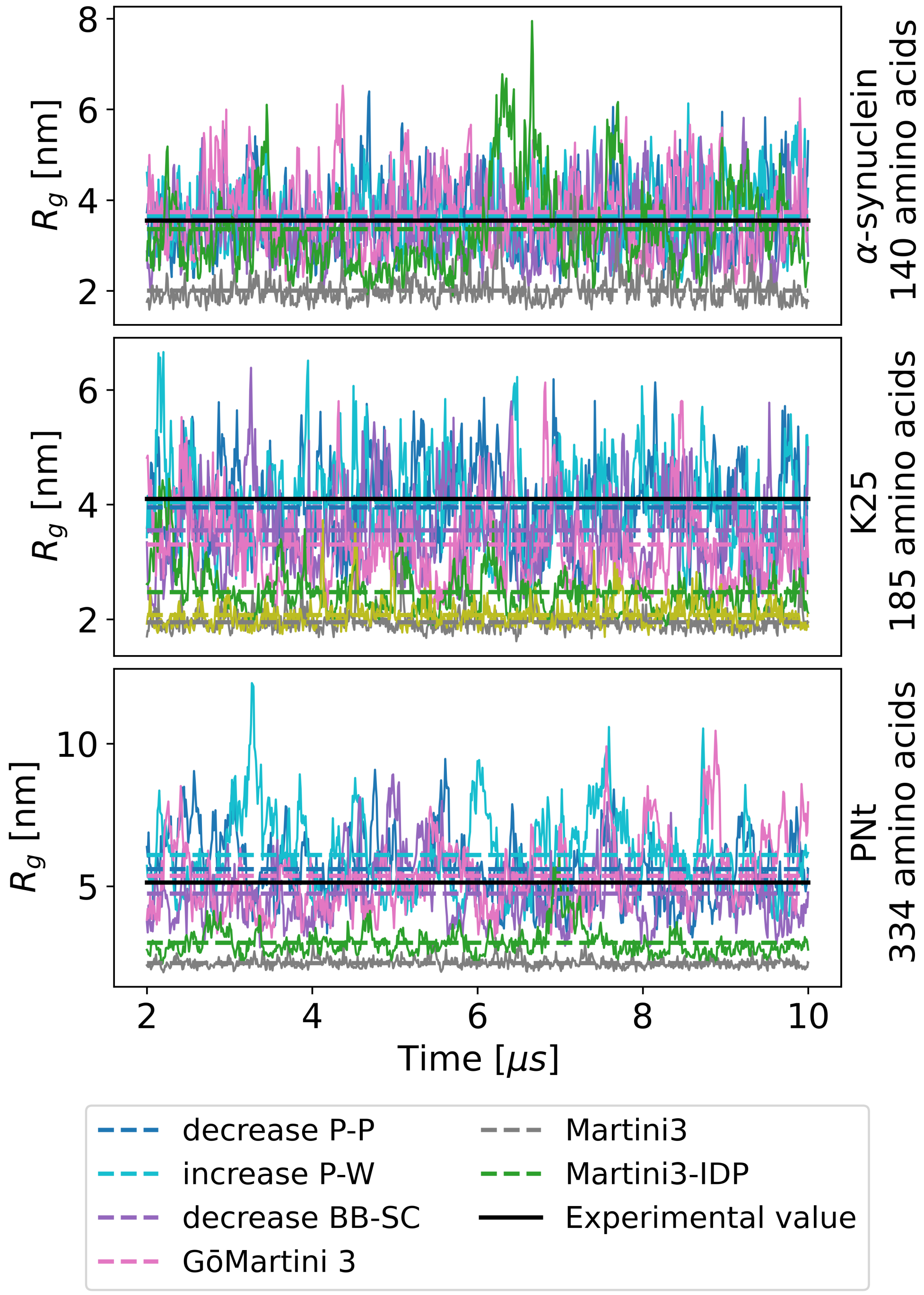


Figure S3: Time traces of the simulation for $\alpha$-synuclein, K25 and PNt. Mean values of $R_g$ are shown as dashed lines for each simulation, and the experimental value as a solid black line.

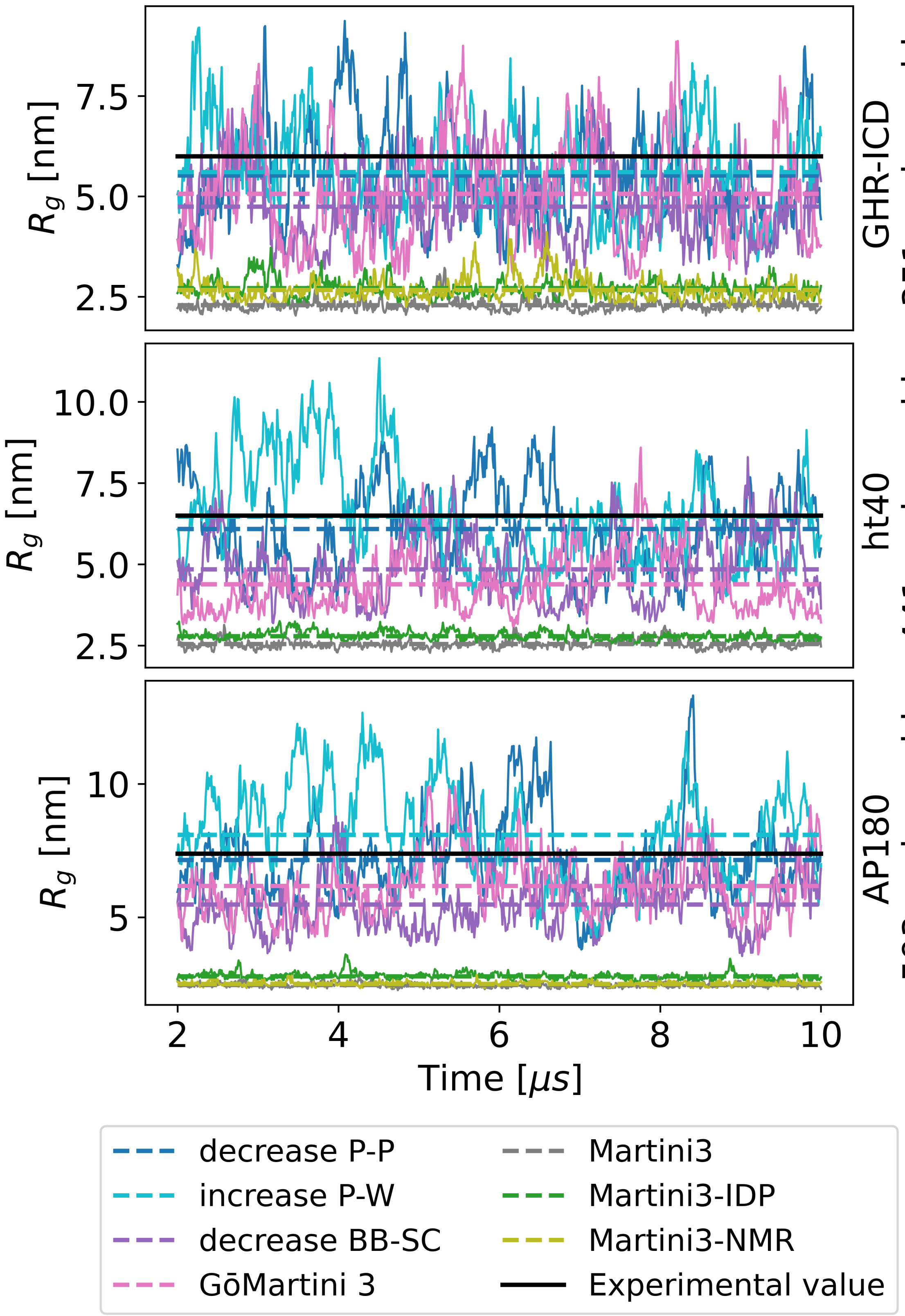


Figure S4: Time traces of the simulation for GHR-ICD, ht40 and AP180. Mean values of $R_g$ are shown as dashed lines for each simulation, and the experimental value as a solid black line.

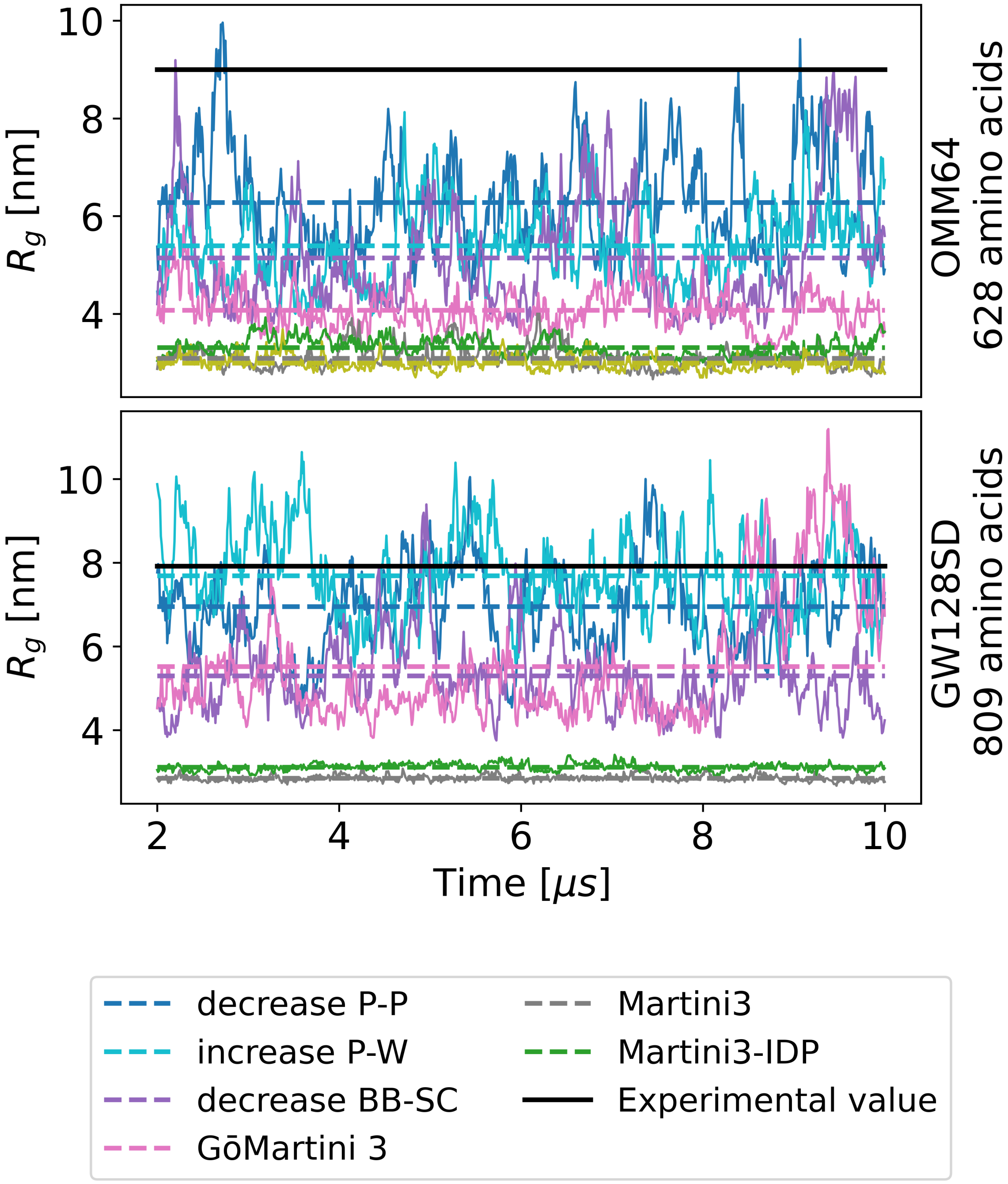


Figure S5: Time traces of the simulation for OMM64 (above) and GW128SD (below). Mean values of $R_g$ are shown as dashed lines for each simulation, and the experimental value as a solid black line.

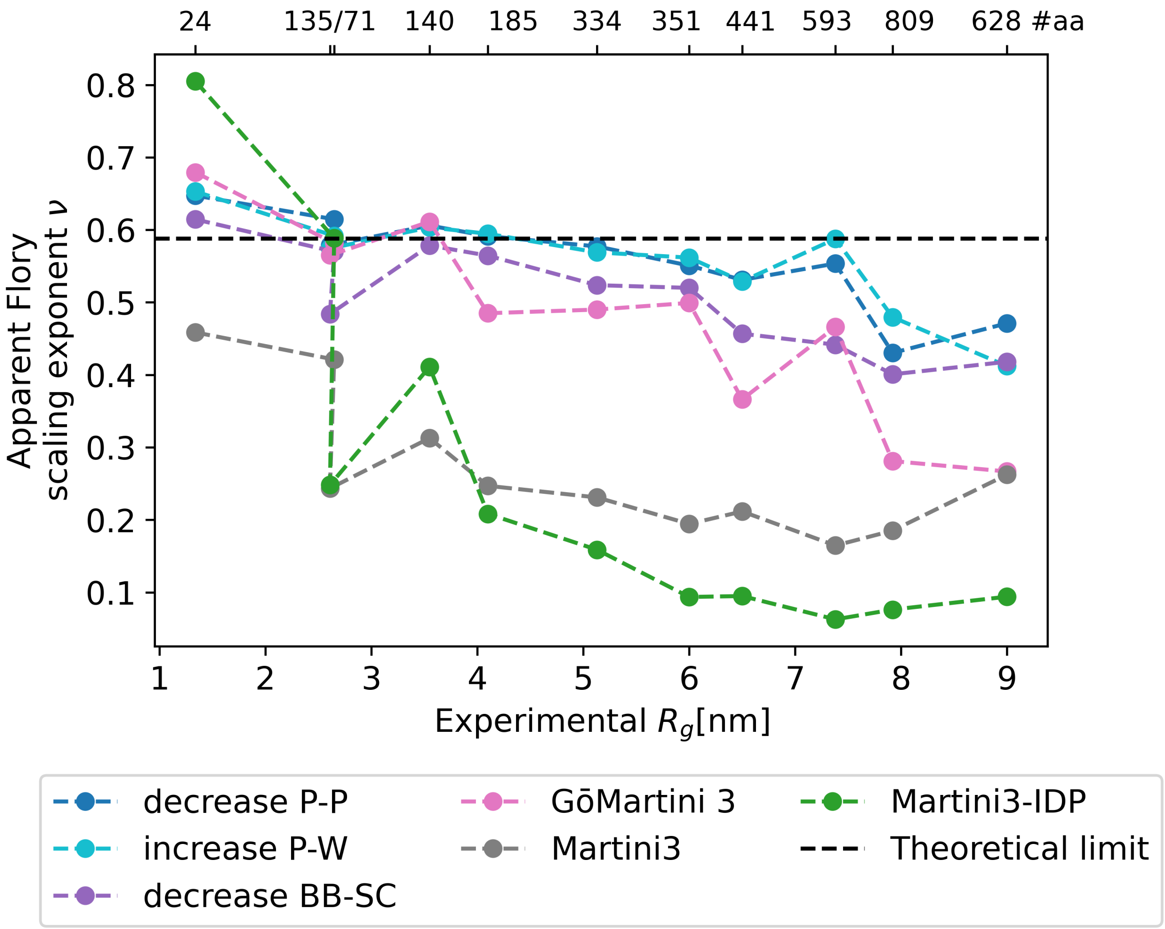


Figure S6: The apparent Flory exponent $\nu$ (that is, it is calculated from a simulation of an IDP) is small for methods which predict compact long IDPs and large for those that predict expanded conformations. The difference in Flory exponents is visible even for small IDPs, before the $R_g$ of the simulated IDP deviates from the experimental $R_g$.

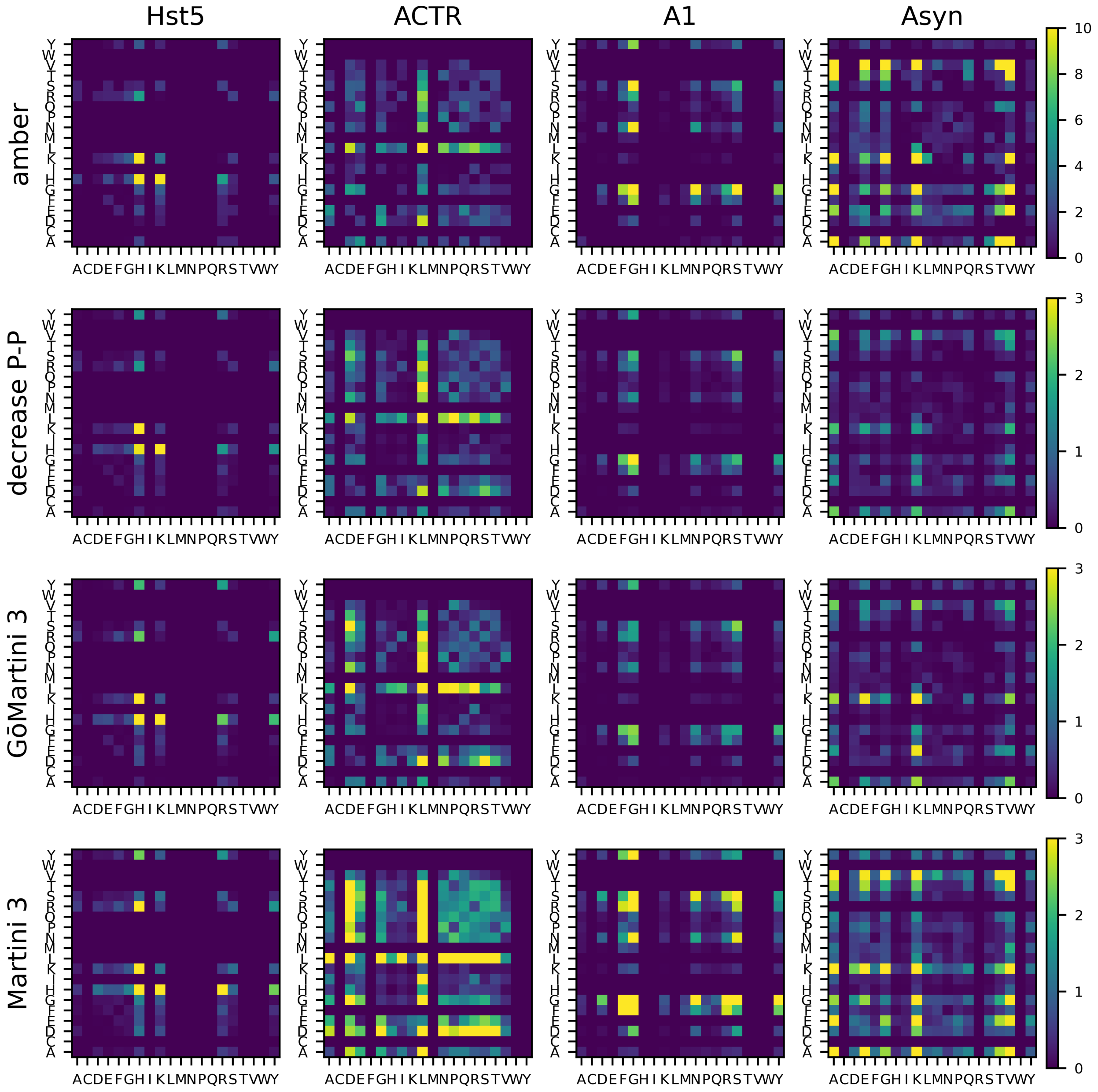


Figure S7: Contact maps by amino-acid comparing the all-atom forcefield amber99-ildn with TIP4P-D water, specialized for IDPs (first row) with the different Martini force fields corrections used in this study (remaining rows). Shown in color is the average number of contacts between the specified amino acid pair per simulation frame. Simulations were performed for four IDPs (columns): Hst5, A1, ACTR and $\alpha$-synuclein. The pattern of which amino-acids preferentially interact with which other amino-acids is very consistent across simulation method, mostly varying in the strength of the contact (number of frames of the simulation in which the amino-acids are in contact).

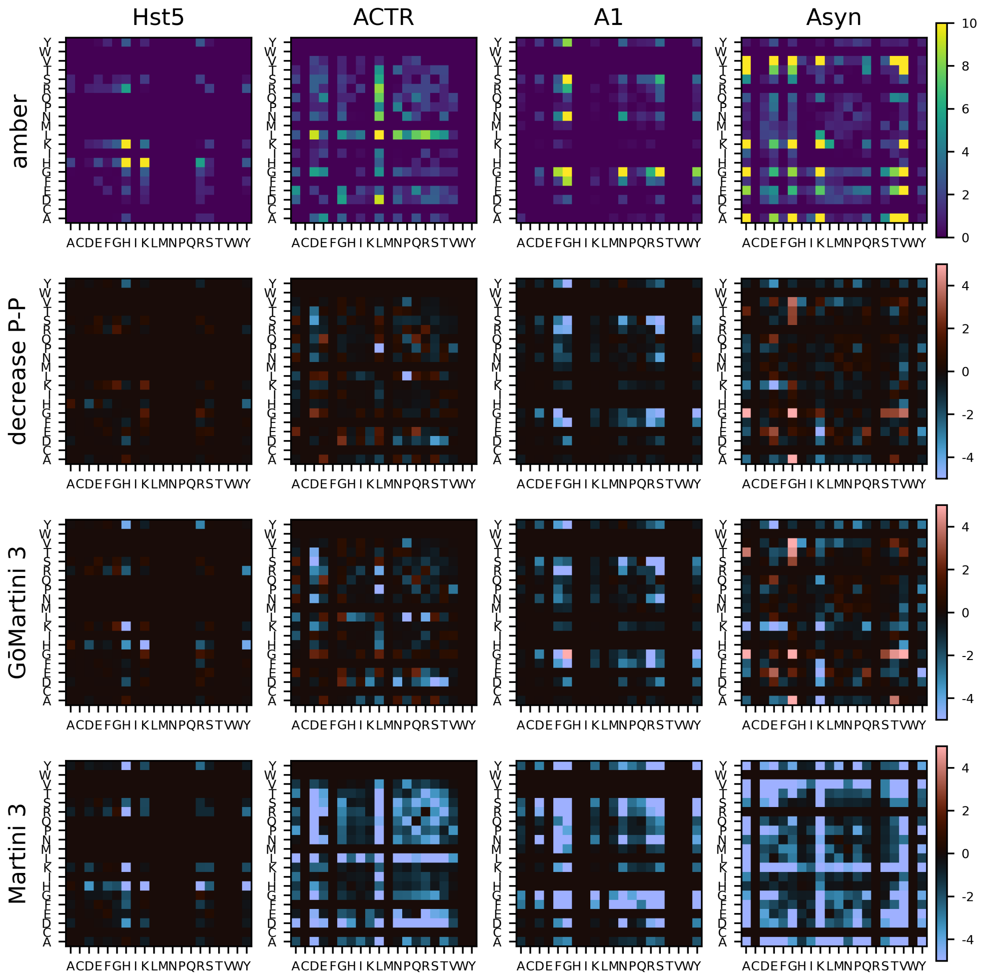


Figure S8: Maps highlighting the difference in contacts between the all-atom forcefield amber99-ildn with TIP4P-D water (specialized for IDPs) (first row) and studied Martini force field corrections (remaining rows). Simulations are performed for four IDPs: Hst5, A1, ACTR and $\alpha$-synuclein (columns). For the all-atom force field, contacts are the average number of contacts between the specified amino acid pair per simulation frame. Contacts for the Martini force-fields are scaled by IDP so that the maximum contacts match for all-atom and Martini with decreased protein-protein interaction and then subtracted from the all-atom maps. Blue indicates that there are more contacts in the Martini field, while red that there are more contacts in the all-atom field.

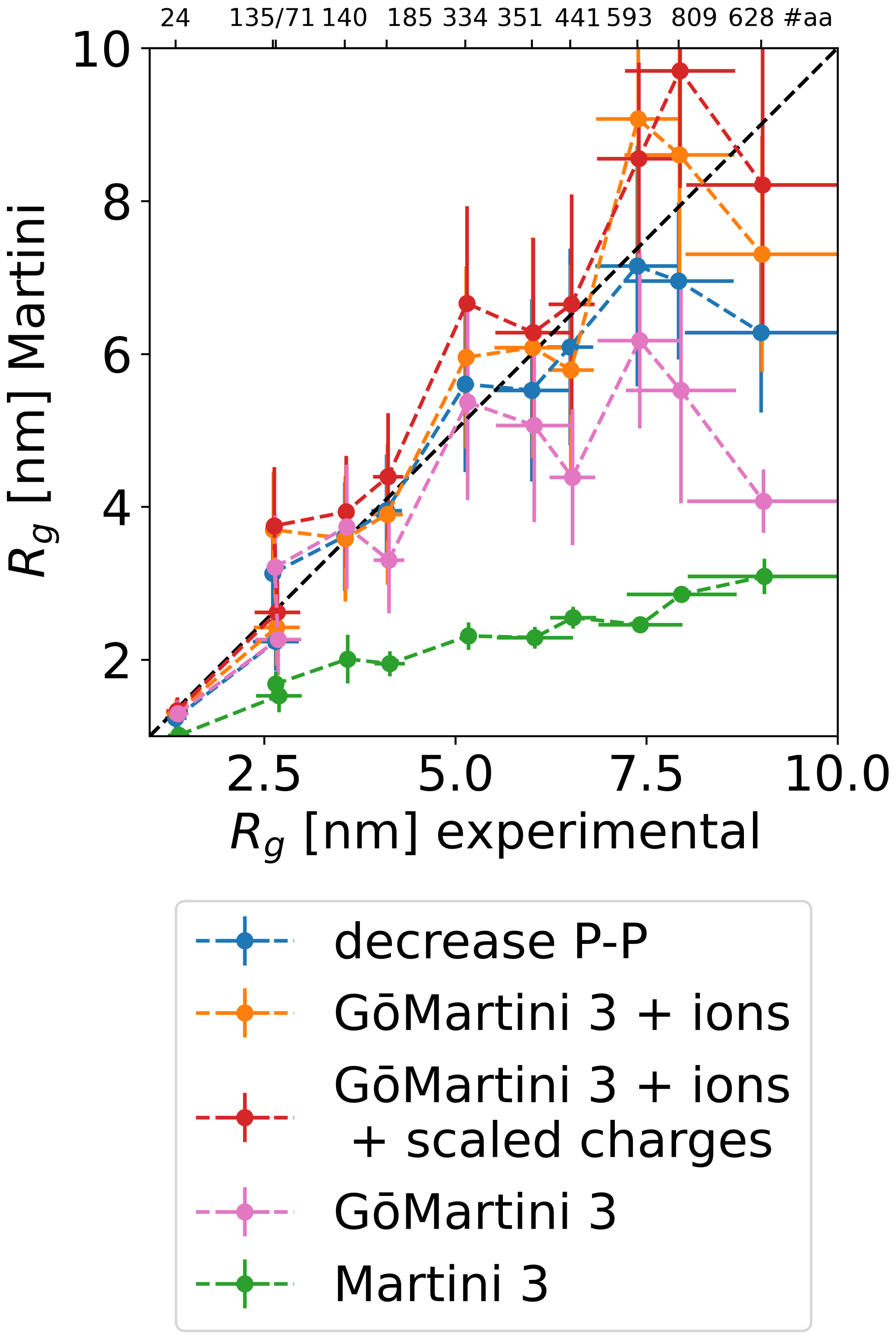


Figure S9: $R_g$ of Martini forcefields using larger and less charged ions (+ ions) or downscaling all charges (+ions + scaled charges) plotted against experimental $R_g$. Martini variants, reducing the protein-protein interactions (decrease P-P) and GōMartini 3, as well as default Martini 3, are included for comparison. The number of amino acids of each studied IDP is indicated at the top.

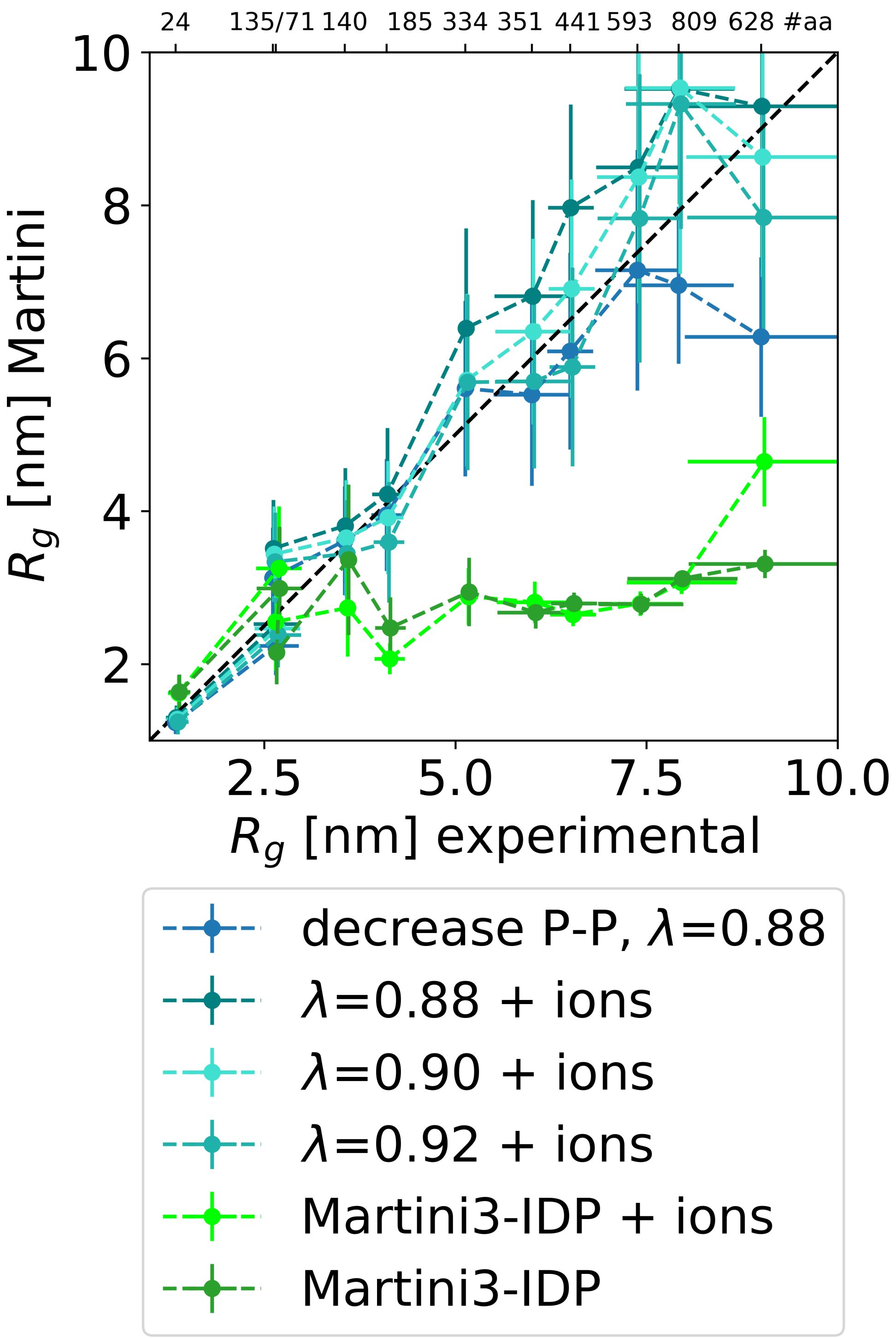


Figure S10: $R_g$ of Martini forcefields where protein-protein interactions have been downscaled as indicated by the factor $\lambda$ in combination with larger and less charged ions (+ ions) is plotted against experimental $R_g$. Martini variants reducing the protein-protein interactions (decrease P-P, $\lambda * 0.88$) or Martini3-IDP are included for comparison. The number of amino acids of each studied IDP is indicated at the top.

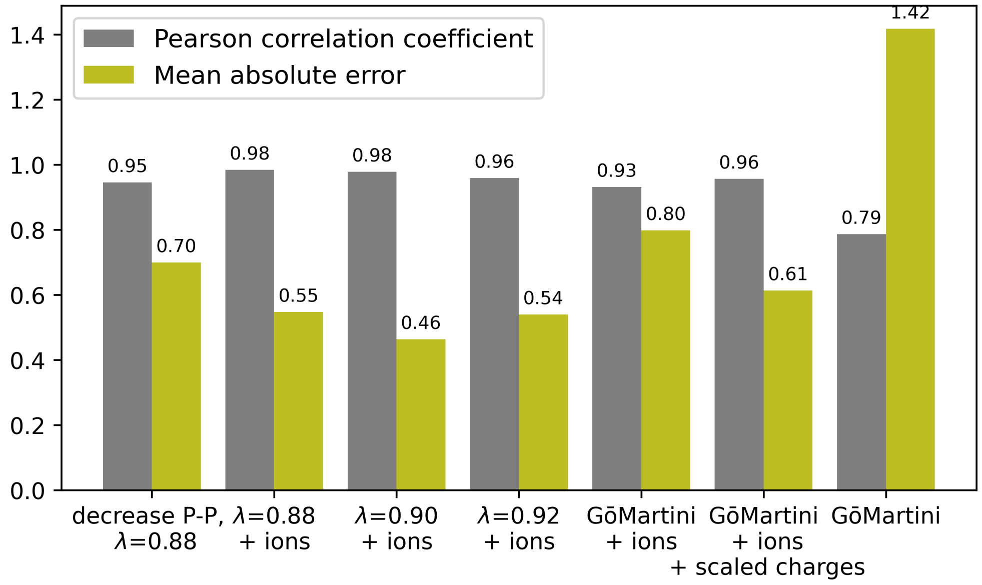


Figure S11: Pearson correlation and mean absolute error on the test set for Martini 3 variations that use larger and less charged ions (see Figures S9, S10)
